# Evidence of reduced viremia, pathogenicity and vector competence in a re-emerging European strain of bluetongue virus serotype 8 in sheep

**DOI:** 10.1101/469460

**Authors:** John Flannery, Beatriz Sanz-Bernardo, Martin Ashby, Hannah Brown, Simon Carpenter, Lyndsay Cooke, Amanda Corla, Lorraine Frost, Simon Gubbins, Hayley Hicks, Mehnaz Qureshi, Paulina Rajko-Nenow, Christopher Sanders, Matthew Tully, Emmanuel Bréard, Corinne Sailleau, Stephan Zientara, Karin Darpel, Carrie Batten

## Abstract

The outbreak of bluetongue virus (BTV) serotype 8 (BTV-8) during 2006-2009 in Europe was the most costly epidemic of the virus in recorded history. In 2015, a BTV-8 strain re-emerged in France which has continued to circulate since then. To examine anecdotal reports of reduced pathogenicity and transmission efficiency, we investigated the infection kinetics of a 2007 UK BTV-8 strain alongside the re-emerging BTV-8 strain isolated from France in 2017. Two groups of eight BTV-naïve British mule sheep were inoculated with 5.75 log_10_TCID_50_ ml^−1^ of either BTV-8 strain. BTV RNA was detected by 2 dpi in both groups with peak viremia occurring between 5-9 dpi. A significantly greater amount of BTV RNA was detected in sheep infected with the 2007 strain (6.0-8.8 log_10_ genome copies mL^−1^) than the re-emerging BTV-8 strain (2.9-7.9 log_10_ genome copies mL^−1^). All infected sheep developed BTV-specific antibodies by 9 dpi. BTV was isolated from 2 dpi to 12 dpi for 2007 BTV-8-inoculated sheep and from 5 to 10 dpi for sheep inoculated with the remerging BTV-8. In *Culicoides sonorensis* feeding on the sheep over the period 7-12 dpi, vector competence was significantly higher for the 2007 strain than the re-emerging strain. Both the proportion of animals showing moderate (as opposed to mild or no) clinical disease (6/8 vs 1/8) and the overall clinical scores (median 5.25 vs 3) were significantly higher in sheep infected with the 2007 strain, compared to those infected with the re-emerging strain. However, one sheep infected with the re-emerging strain was euthanized at 16 dpi having developed severe lameness. This highlights the potential of the re-emerging BTV-8 to still cause illness in naïve ruminants with concurrent costs to the livestock industry.

**Summary:** The re-emerging Bluetongue virus serotype 8 still presents a threat to naïve ruminants in Europe despite reduced virulence

## Introduction

Bluetongue (BT) is an infectious hemorrhagic disease of ruminants caused by bluetongue virus (BTV) which is transmitted via *Culicoides* biting midges (Carpenter et al., 2013). BTV is the type species of the genus *Orbivirus* within the family *Reoviridae* and is a serologically and genetically diverse virus. The BTV genome is comprised of 10 double-stranded RNA segments encoding several structural and non-structural proteins. BTV segment-2 encodes the most variable BTV protein (VP2) which contains the majority of epitopes that interact with neutralising antibodies (Maclachlan et al., 2014). Since 1998, a number of BTV serotypes have caused both sporadic and widespread incursions into the EU (Belbis et al., 2017).

In August 2006, a BTV serotype 8 (BTV-8) strain of sub-Saharan origin (Maan et al., 2008) was detected within animal holdings in The Netherlands, the first time that the virus had been identified in northern Europe. This BTV-8 strain successfully re-emerged in 2007 and subsequently spread throughout most northern European countries, causing widespread clinical disease and major economic damage to the farming sector (Wilson and Mellor, 2009). However, in conjunction with animal movement restrictions, the implementation of vaccination campaigns by EU member states was effective in reducing and eventually preventing further transmission of BTV-8 and only sporadic outbreaks of disease occurred in the EU by 2009.

In August 2015, BTV-8 re-emerged in France and subsequently spread throughout the entire country (Sailleau et al., 2017). How BTV-8 persisted in areas that were thought to have been free of virus transmission remains unknown, although it has been recently proposed that low-level circulation of BTV-8 occurred in France prior to the detection in 2015 (Courtejoie et al., 2018). Importantly however, anecdotal observation suggested that changes had occurred in the epidemiology of the re-emerging strain. Whereas the 2007-2009 BTV-8 strain caused widespread clinical signs in cattle and sheep (Elbers et al., 2009, Zanella et al., 2013), the re-emerging BTV-8 strain has thus far only caused mild clinical illness (Sailleau et al., 2017). In addition, the rate of spread of the virus in France appeared slower than the original BTV-8 strain. At the time of the BTV-8 re-emergence in 2015, it has been estimated that the ruminant herd immunity in France was 18% (Bournez et al., 2018).

Genetically, the re-emerging BTV-8 strain (GenBank accession numbers: KP56990-KP56999) differs from a 2007 UK BTV-8 strain (accession numbers: KP820957, KP821077, KP821199, KP821319, KP821439, KP7821559, KP821681, KP821801, KP821921 and KP822042) in just 11 amino acids occurring in segments 1 (3), 2 (1), 3 (1), 4 (1), 8 (1), 9 (3) and 10 (1). The high degree of amino acid similarity between the 2007 and 2015 strains underpins the hypothesis that this is not a new introduction of BTV-8 (Sailleau et al., 2017).

In this study we directly compare the infection of sheep with the original and re-emerging European BTV-8 strains by assessing viremia, clinical signs, antibody production and transmission to *Culicoides* biting midges. By using British mule sheep, representative of the UK flock from a region not originally exposed to BTV-8 infection, the aim is to understand if the changes in epidemiology and pathogenicity observed in France will also occur in regions beyond those affected by outbreaks in 2006-2009. The findings from this study will inform response in both the UK and other countries previously unaffected by BTV-8 and allow for evidence-based surveillance and control measures to be implemented.

## Materials and methods

### Ethical statement for animal studies

All animal experiments were carried out in accordance with the UK Animal Scientific Procedure Act (ASPA) 1986 which transposes European Directive 2010/63/EU into UK national law. The animal studies were approved by the UK Home Office in granting Project licence 70/7819 under the Animal Scientific Procedure Act and all protocols underwent appropriate local ethical review procedures by the Animal Welfare and Ethical Review Board (AWERB) of The Pirbright Institute.

### Preparation of inocula

A 2017 French BTV-8 strain (FRA2017) which had been isolated on *Culicoides sonorensis* (KC) cells through a single passage was provided by the French BTV NRL (Animal Health Laboratory, ANSES, Maisons-Alfort). BTV-8 isolate UKG2007/03 (UKG2007), isolated through a single passage from the 2007 UK index case was obtained from the Orbivirus Reference Collection, (Pirbright Institute, UK). Both the UKG2007 and FRA2017 strains were re-passaged once on KC cells. Following harvest, viral inocula were titrated on KC cells using a 96-well titration assay in combination with immunofluorescence microscopy. Briefly, 10-fold serial dilutions of the virus were added to KC cells immediately following addition of the cells to 96 well tissue culture plates. After 5 days incubation at 26°C, the cells were fixed using 4% paraformaldehyde for 45 min and following multiple washes with PBS were permeabilized with 0.2% Triton X-100 for 20 min following. Viral proteins were labelled using an anti-BTV polyclonal guinea-pig sera (ORAB279) and anti-guinea-pig-Alexa488 and immunofluorescence was visualized on a Nikon Eclipse TE300 microscope to calculate viral titers according to Spearman-Kärber. Subsequently, the inocula were adjusted using Schneiders media to yield a final titer of 5.75 log_10_ TCID50/mL.

### Study design

Eighteen adult female British mule sheep, representing the most common British mixed breed of > 7 years of age were chosen for use in this study. All sheep were previously tested as BTV-antibody negative by C-ELISA. The sheep were randomly assigned (see supplementary information) into two groups of 9 individuals. The two groups were housed in separate rooms of the Pirbright Institute’s high containment animal facility. Eight sheep per group were inoculated with 1 mL subcutaneously in the left neck and 0.5 mL intradermally distributed over 5 inoculation sites into the inner left thigh of either UKG2007 or FRA2017. One sheep in each respective group was kept uninfected as a contact transmission control (sheep 2 and sheep 18). Blood samples (EDTA and whole blood) were taken from the jugular vein at −1, 2, 3, 5, 6, 7, 8, 9, 10, 12, 14, 16, 19, and 20/21 dpi. Sheep body temperatures were recorded daily and clinical scoring was performed throughout as described previously (Darpel et al., 2007).

### Vector competence

*Culicoides sonorensis* from a colony maintained at The Pirbright Institute (PIRB-s-3 strain) were fed on two sheep from each group at 6 dpi (UKG2007) or 7 dpi (FRA2017) to coincide with determined peak viremia as described previously (Baylis et al., 2008). Engorged individuals were incubated at 25°C for 8 days and surviving individuals were homogenized in GMEM using a tissue lyser (Qiagen) and made up to a final volume of 1 mL per sample (Veronesi et al., 2013). *Culicoides* samples were analyzed in pools of eight individuals for BTV genome detection by RT-qPCR as described below. Pools that reported a cycle threshold (C_T_ value) less than that of the original blood meal in midges harvested on the day of feeding were considered to contain one or more individuals competent for BTV transmission.

## Molecular analyses

### Preparation of plasmid for quantitation in real-time RT-PCR assay

Plasmid pGEM-3Zf(+) carrying a 97 bp sequence from BTV segment 10 (derived from the assay described by (Hofmann et al., 2008)) was transformed into JM109 competent cells (Promega), purified and then quantified using a nanodrop to create working stocks of 10^6^ BTV genome copies/μL.

### RNA extraction and RT-qPCR analysis

BTV RNA was extracted from 100 μL of EDTA blood and eluted into 80 μl buffer using the KingFisher Flex automated extraction platform and the MagVet Universal nucleic acid extraction kit (ThermoFisher Scientific, Paisley, UK). Ten microliters of sample RNA was analyzed as per the assay described by (Hofmann et al., 2008) with modifications (Flannery et al., 2018) using the Express One-Step qRT-PCR kit (ThermoFisher) on an Applied Biosystems 7500 Fast instrument (ThermoFisher). A log-dilution series of the plasmid (1 × 10^0^ to 1 × 10^6^ copies per μL) was included in triplicate on each RT-qPCR run. BTV RNA copies were determined by comparing sample C_T_ values to the standard curves and expressed as BTV genome copies/mL.

### Serological analyses

Whole blood samples were centrifuged at 3000 × g for 5 min and serum was decanted and stored at +4°C until analysis. BTV antibodies were detected using the ID Screen^®^ Bluetongue Competition kit (ID Vet, Grabels, France) in accordance with the manufacturer’s instructions. The serum neutralisation test (SNT) against BTV-8 was performed as described previously (Batten et al., 2012).

### Virus isolation

EDTA blood cells were washed 3× with PBS and sonicated as described in the World Organisation for Animal Health (OIE) manual (Savini, 2014). KC cells were inoculated with 100 μl of washed blood and incubated at 26°C in Schneiders media (1% Amphotericin B, 1% Penicillin/Streptomycin and 10% FBS). Media was replenished 24 hours post infection and cells were incubated for a further 6 days. Following the 7-day incubation, cells were harvested, centrifuged at 3000 × g for 5 min and the supernatant tested for BTV using RT-qPCR as described above. Virus was considered to have been isolated if the BTV C_T_ value of the harvested material was 3-C_T_ values less than that of the original EDTA-blood inoculum.

### Statistical analysis

Body temperatures and clinical scores were analyzed by comparing the maximum values and the dpi at which they occurred for each strain using Wilcoxon rank-sum (also known as Mann-Whitney U) tests. The proportion of sheep showing moderate (as opposed to mild or no) clinical disease in each group was compared using a Fisher exact test. Viremia for sheep infected with each strain was analyzed by comparing the maximum titer, the dpi at which it occurred and the area under the curve (a measure of total virus production; computed using the trapezium rule) using Wilcoxon rank-sum tests. Non-parametric tests were preferred because of the small group sizes and potential non-normality of the data.

The vector competence for each BTV-8 strain was calculated from the number of positive pools and the number of pools tested using a binomial likelihood and noting that the probability of a positive pool is equal to 1-(1-*b*)^*n*^, where *b* is the vector competence (i.e. the probability of an individual midge becoming infected) and *n* is the number of midges in the pool. A second model was also considered in which vector competence was independent of strain, but depended on the viral titer (log_10_ genome copies/mL) in the sheep on which each pool of midges fed, so that *b*=1/(1+exp(−*a*_0_−*a*_1_×titer)).

## Results

### Clinical observations

Typical clinical signs of BTV infection (fever, depression, facial edema, reddening of the mucosal membrane and coronary bands) were seen more frequently and were more pronounced in the UKG2007-inoculated sheep compared to the FRA2017-inoculated sheep (figure 1). Overall, 6/8 UKG2007-inoculated sheep were classed as moderately ill, while two sheep were classed as mildly ill. By contrast, only one sheep was classed as moderately affected in the FRA2017-inoculated sheep, with the remaining 7/8 being classed as mildly affected. The proportion of sheep showing moderate (as opposed to mild) clinical disease was significantly (p = 0.04) higher in sheep infected with UKG2007 compared with those infected with FRA2017. In addition, the maximum clinical score was significantly (p = 0.014) higher in sheep infected with UKG2007 (median score 5.25, range 2-6) compared with those infected with FRA2017 (median score 3, range 1.5-3.5), but the time at which the maximum score occurred did not differ significantly (p = 1.0) between the two groups of BTV-8 inoculated sheep.

**Figure 1.**
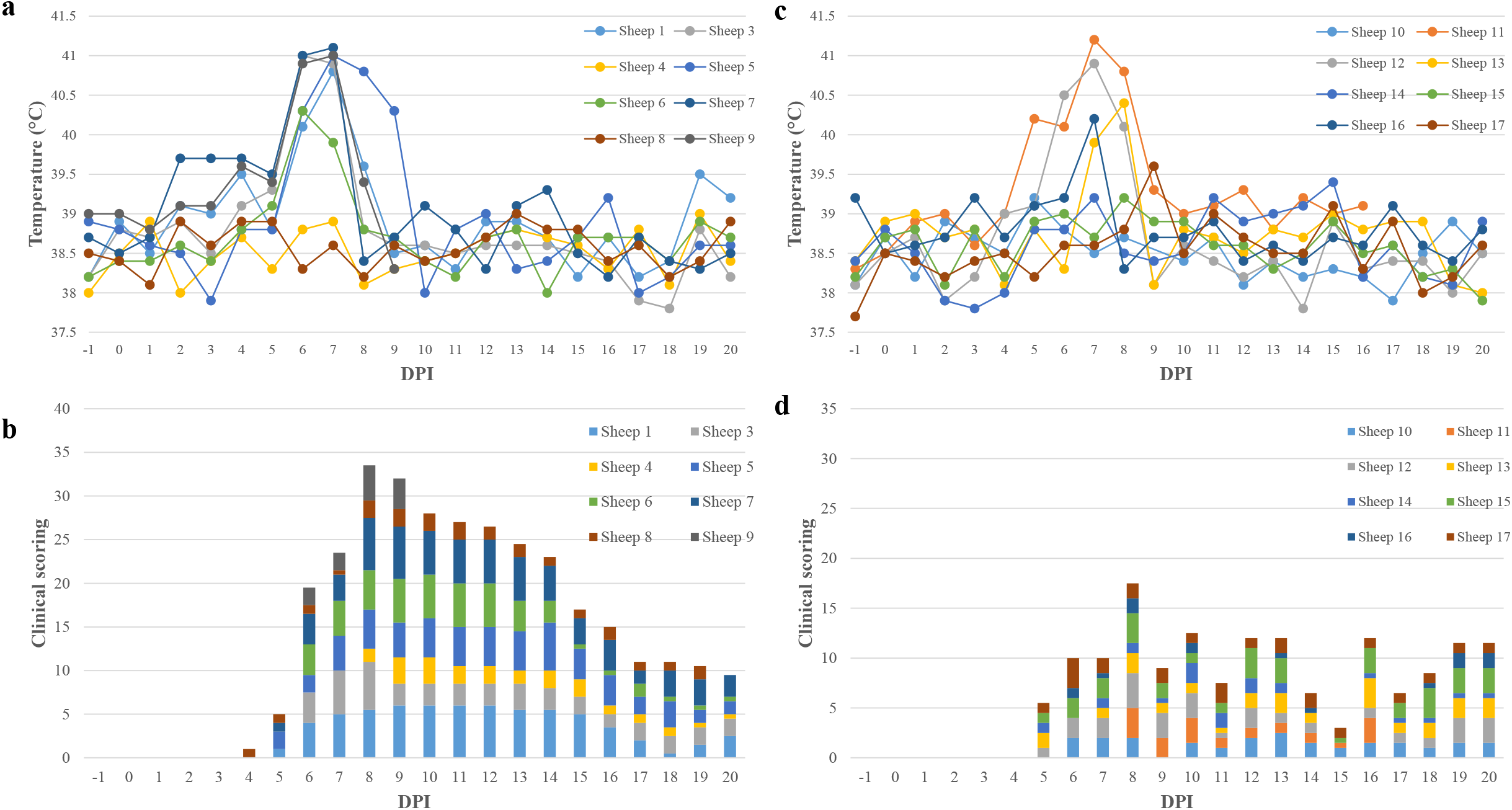
Daily temperatures of sheep inoculated with UKG2007 (a) and FRA2017 (c), and combined clinical scores of sheep inoculated with UKG2007 (b) and FRA2017 (d).

Five of eight UKG2007-inoculated sheep showed pyrexia (>40.5°C), in contrast with 2/8 FRA2017-inoculated sheep (figure 1). However, the difference between the groups was not statistically significant for either maximum temperature (p = 0.49) or the dpi at which this occurred (p = 0.76).

Two sheep were euthanized on welfare grounds during the study, one in each group. The sheep from the UKG2007-inoculated group was euthanized at 9 dpi for reaching the humane endpoint (at the end of the moderate clinical spectrum) of the moderate study protocol. In this sheep, typical hemorrhagic lesions on mucosal membranes, coronary band, lymph nodes, pulmonary artery as well as the epithelium of the reticulum and omasum were observed during necropsy. The sheep from the FRA2017-inoculated group developed acute lameness on all four feet late in infection and was euthanized on humane grounds at 17 dpi.

The two transmission control sheep remained clinically unaffected other than exhibiting a mild nasal discharge over a two-day period during the experiment.

### Molecular analyses

Figure 2 shows the BTV concentrations in EDTA blood over the study period. In 5 UKG2007-inoculated sheep and in 3 FRA2017-inoculated sheep BTV RNA was detected by RT-qPCR at 2 dpi. BTV RNA was detected in all other inoculated sheep at 5 dpi with the exception of sheep 17 (that had been inoculated with FRA2017). Peak viremia occurred earlier for UKG2007-inoculated sheep (between 5 and 6 dpi) than for FRA2017-inoculated sheep (5-9 dpi), but this difference was not statistically significant (p = 0.11). At peak viremia, BTV concentrations in blood (log_10_ genome copies/mL) were significantly (p = 0.021) higher for UKG2007-inoculated sheep (median 7.0; range: 5.5-8.3) compared with FRA2017-inoculated sheep (median 5.4; range: 1.6-7.3). Furthermore, the area under the curve (a measure of total virus production) was significantly (p = 0.028) greater for UKG2007-inoculated sheep (median 7.5; range: 6.0-8.8) compared with FRA2017-inoculated sheep (median 5.9; range: 2.9-7.9).

**Figure 2.**
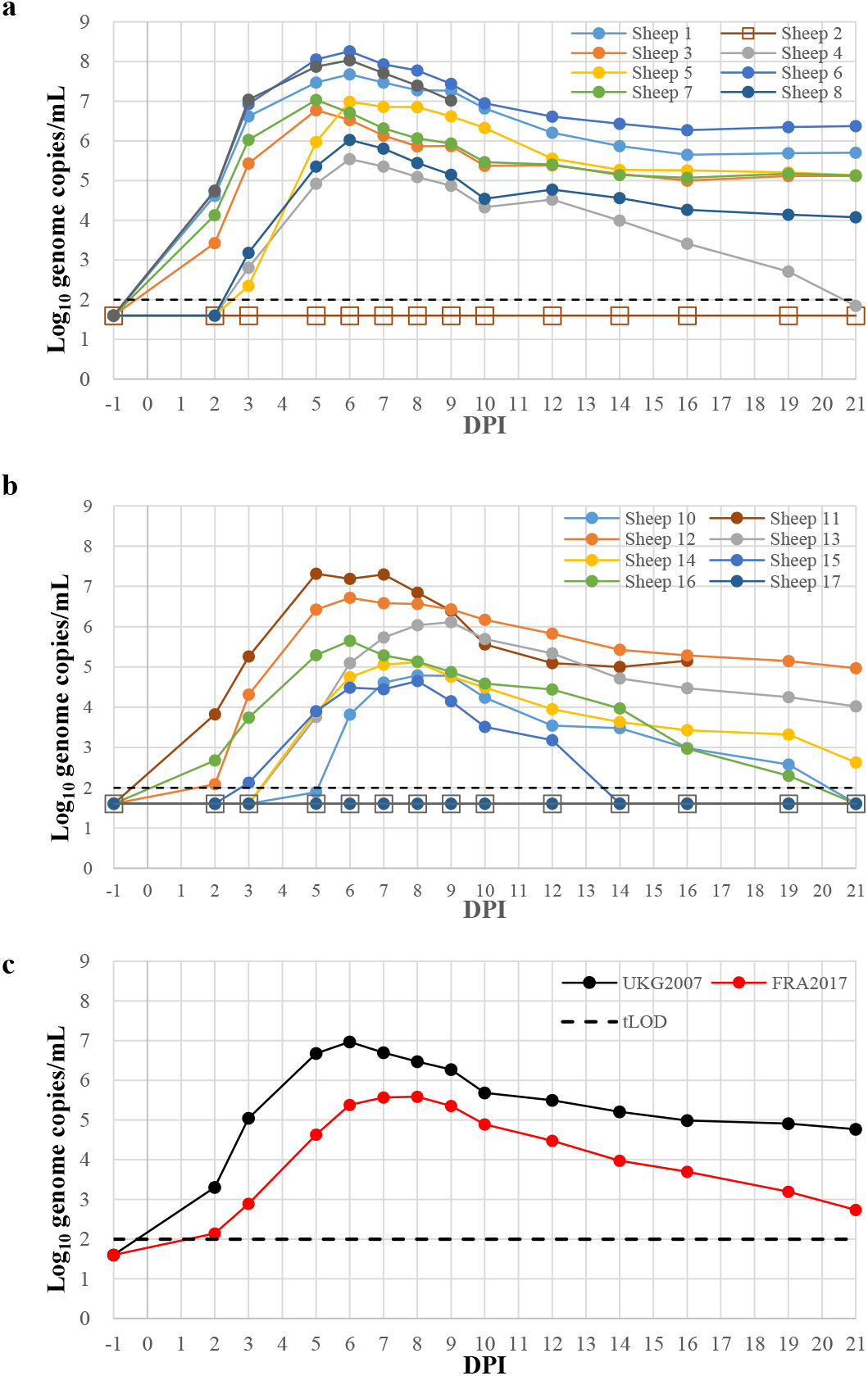
BTV genome concentrations in sheep inoculated with UKG2007 (a) and FRA2017 (b) and mean genome concentrations for both groups (c). Sheep 2 and Sheep 18 are the control animals.

At 21 dpi in the remaining animals, 3 sheep inoculated with the FRA2017 strain had cleared the infection. BTV remained detectable in all 7 sheep and in 3 sheep inoculated with UKG2007 and FRA2017 strains, respectively. At the end of the experiment (21 dpi), mean BTV concentrations were 5.26 (range: 4.08-5.70) and 3.87 (range: 2.63 and 4.97) log_10_ genome copies/mL for UKG2007 and FRA2017-inoculated sheep, respectively. BTV RNA was not detected in the transmission controls or in sheep 17 throughout the experiment.

### Vector competence

A total of 272 individual *C. sonorensis* fed on sheep 6 and sheep 9 that were infected with the UKG2007 strain survived incubation and this equated to 34 pools of midges for testing. A total of 280 individuals fed on sheep 11 and sheep 12 that were infected with FRA2017 were tested in 36 pools. Three midge pools from the UKG2007-inoculated sheep had lower C_T_ values than the mean value obtained in *C. sonorensis* tested on the day of feeding (C_T_ 30.9). *C. sonorensis* tested on the day of feeding on FRA2017-infected sheep had mean C_T_ value of 34.7. BTV RNA was not detected in any midge pools from the FRA2017-inoculated sheep.

Based on these results for pooled midges, the estimated vector competence (i.e. probability of an individual midge becoming infected) was 1.2% (95% confidence interval (CI): 0.3% to 3.0%) for UKG2007 and 0% (95% CI: 0% to 0.7%) for FRA2017, which are significantly different. Viral titers (log_10_ genome copies/mL) in the sheep infected with UKG2007 were higher on the day of *Culicoides* feeding than for those infected with FRA2017 (8.3 and 8.0 compared with 7.3 and 6.6, respectively). An increase in competence was associated with an increase in titer, but this was not statistically significant (estimate for coefficient, *a*_1_: 2.6; 95% CI: −0.1 to 3.6).

### Serological analyses

Figure 3 shows the BTV antibody response using C-ELISA. In the sheep inoculated with UKG2007, five of the eight seroconverted by 7 dpi, and by 8 dpi all inoculated sheep had seroconverted (defined as <40% inhibition by the manufacturer). In the group inoculated with FRA2017, none of the sheep seroconverted at 7 dpi yet all had seroconverted by 9 dpi. Based on the ELISA results, SNTs were performed on serum samples obtained at 7, 9, 12, 14 and 21 dpi. Neutralizing antibodies were first detected at 7 and 9 dpi for UKG2007 and FRA2017-inoculated sheep, respectively. By 14 dpi, all sheep had neutralizing antibodies with titers (log_10_) between 1.18 and 2.68, rising to between 1.60 and 2.81 at 21 dpi. The mean antibody titers at 21 dpi were 2.01 and 1.54 for the UKG2007 and FRA2017-inoculated sheep, respectively. The transmission control sheep in both groups did not seroconvert.

**Figure 3.**
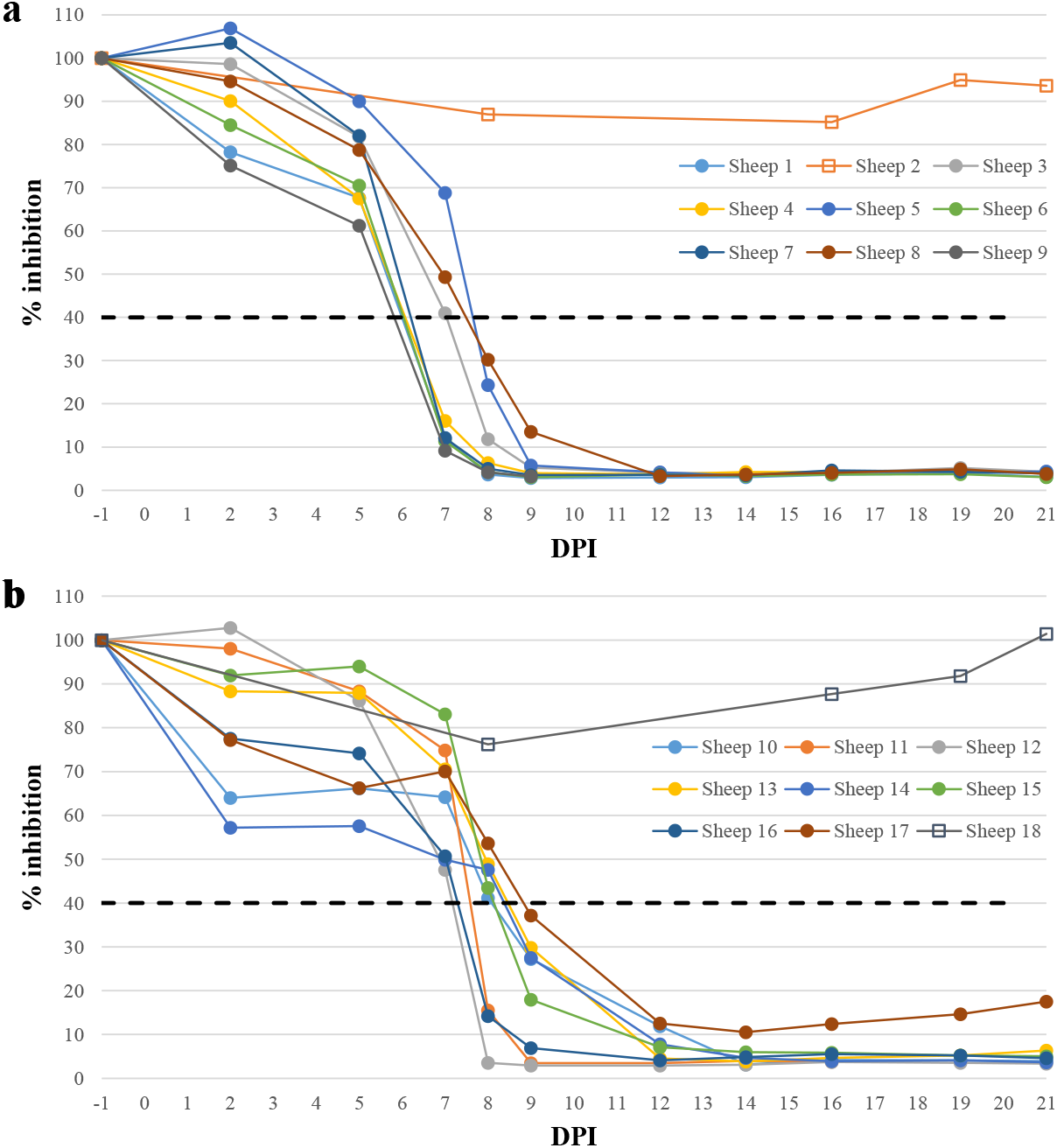
Antibody response of sheep inoculated with UKG2007 (a) and FRA2017 (b) as determined by C-ELISA. The dashed line represents the positive/negative cut off of the ELISA whereby values <40% inhibition are considered positive. Sheep 2 and Sheep 18 are the control animals.

### Virus isolation

Virus isolation from EDTA blood was performed for all inoculated sheep at 2, 5, 7, 10, 12, 14, 16 and 21 dpi. For UKG2007-inoculated sheep, virus was isolated at 2 dpi (3/8 sheep), 5 dpi (7/8 sheep), 7-10 dpi (8/8 sheep) until 12 dpi (1 sheep). For FRA2017-inoculated sheep, virus was isolated at 5-7 dpi (6/8 sheep) until 10 dpi (4/8 sheep).

## Discussion

Constant and largely unpredictable outbreaks of BTV present a significant global challenge in terms of proportionate response. This is exacerbated by our inability to accurately define the pathogenicity of emerging and re-emerging BTV strains, which is the main determining factor in vaccine production by industry and subsequent uptake by farmers. The aim of this study was to compare infection kinetics and clinical severity of a re-emerging BTV-8 strain in sheep to understand the impact of this strain in areas not affected by BTV-8 during 2006-2009. The study confirms anecdotal evidence from the field that viremia and clinical signs are reduced in the re-emerging strain. However, this strain may significantly impact upon naïve sheep populations as evidenced by one sheep developing acute lameness during the convalescence phase of infection.

British mule sheep were used in this study as representative of the UK flock and seem slightly more resistant to the UKG2007 strain than pure-bred Dorset Poll sheep when infected with the UK BTV-8 strain (Moulin et al., 2012). Indeed, one sheep inoculated with the 2007 strain had almost cleared the infection by the end of the experiment (21 dpi). This variability in the severity of BTV infection has been attributed to numerous host factors, such as the breed, health and age of the ruminant (Maclachlan et al., 2009, Caporale et al., 2014). It has been previously reported that in field settings, environmental factors, such as exposure to sunlight, elevated temperatures, stress and bacterial or viral co-infections can exacerbate BT clinical signs in sheep (Kyriakis et al., 2015). Therefore, the clinical presentation of infection with the BTV-8 strains in the field is likely to be more severe than what we have reported in this study where animals were housed within the high-sanitary and controlled-husbandry conditions of the containment facilities. This highlights that infection with the re-emerging BTV-8 strain may still have a considerable economic impact in the field.

Interestingly, one of the sheep inoculated with the re-emerging BTV-8 strain did not develop detectable viremia throughout the entire study, yet seroconverted and developed a neutralizing antibody response. This suggests that the sheep did receive the inoculum. Historically studies have previously reported that ruminants experimentally-inoculated with BTV did not develop detectable viremia (in the presence or absence of seroconversion), although viremia detection was based on virus isolation rather than the more sensitive use of real-time PCR (Parsonson et al., 1987, Flanagan et al., 1982, Baylis et al., 2008). Other studies have further noted seroconversion of individual BTV-inoculated sheep or cattle either in the presence of very low detectable RNA levels by RT-qPCR (van der Sluijs et al., 2013) or in the absence of detectable viremia, however, the latter was associated with either a low-titer (Baylis et al., 2008, Di Gialleonardo et al., 2011) or a high-passage inoculum (Janowicz et al., 2015). Interestingly, a recent study also highlighted that a highly-passaged BTV-8 strain infected and was detectable in the skin and draining lymph nodes of inoculated sheep however did not result in detectable RNA levels in the systemic blood (Melzi et al., 2016). Overall all these studies, including the data presented here, demonstrate that manifestation of clinical disease and viremia varies in individual animals even when infected by the same route and with the same dosage of virus, highlighting differences in individual susceptibility to BTV infection. Alternatively, it is possible that this sheep had residual immunological memory from previous vaccination which could provide some protection from infection even in the absence of detectable antibodies. BTV antibodies can be detected in sheep up to 2.5 years following vaccination (Batten et al., 2013) and it has been estimated that they can persist for 5-6 years following infection or vaccination (Bournez et al., 2018). The sheep used in our study were at an advanced age (>7 years old), as reported by the farmer and by assessing the general condition of the sheep.

Significantly greater RNA copy numbers were detected in sheep inoculated with the 2007 strain than the re-emerging strain. In addition, BTV was isolated more frequently from sheep inoculated with the 2007 strain. By the end of the experiment, 3 sheep inoculated with the re-emerging strain had also cleared infection (BTV RNA was not detected). This reduced viremia was reflected in the quantity of BTV RNA imbibed by *C. sonorensis* implying that monitoring through the venous route was representative of the quantity of viral RNA in the skin. Furthermore, *C. sonorensis* demonstrated a higher vector competence (1.2%) towards the 2007 strain than the re-emerging strain. The lower vector competence observed using FRA2017 is possibly related to lower viral titers in the blood of sheep infected with this strain, though the small number of sheep used for the vector infection studies preclude drawing robust conclusions about the relationship between competence and viremia. Overall, these results suggest that the re-emerging strain elicits a lower viremia, for a shorter duration in sheep and is less efficient at infecting *Culicoide*s biting midges. These findings are consistent with the relatively slow spread of the re-emerged BTV-8 strain which has remained largely within the French borders since circulation was detected in August 2015.

The infection kinetics of the two BTV-8 strains were similar to each other and were in-line with those reported for other BTV serotypes in sheep (Schulz et al., 2018, van der Sluijs et al., 2013). However, the re-emerging strain was found to be less virulent than the 2007 strain. It has been postulated that the re-emerging strain has been circulating asymptomatically in wildlife since the end of the 2006-2009 BTV-8 epizootic (Sailleau et al., 2017). The reason behind the attenuation of BTV strains over a period of time has been attributed to selective pressures placed on circulating BTV serotypes during replication within their mammalian hosts and vectors (Caporale et al., 2014). The respective lineages of the two BTV-8 strains used in this study differ by a few amino acids, thus, the impact of future genomic mutations in the re-emerging BTV-8 strain is uncertain. Nonetheless, detailed genomic comparison between different BTV strains used in experimental infection studies may increase our overall understanding of the molecular determinants of BTV virulence.

We have performed this experimental infection study in sheep; therefore, the impact of the re-emerged BTV-8 strain in cattle is yet to be determined experimentally. Previous experimental infection studies of other European BTV-8 strains in cattle did not result in overt disease or caused mild clinical signs which were in contrast with the clinical illness reported in sheep (Darpel et al., 2007, Martinelle et al., 2011, Di Gialleonardo et al., 2011). An experimental comparison of BTV-8 strain-specific virulence (similar to our study here) is likely to be more challenging in cattle.

Considering that morbidity rates in field conditions are likely to be higher than what we have reported here and coupled with animal movement restrictions, it is probable that future incursions of the re-emerged BTV-8 will have an economic impact on the livestock industry. Therefore, appropriate surveillance and vaccination strategies should be considered in advance of the likely incursion of this re-emerged BTV-8 strain.

## Acknowledgements

This study was funded by the European Commission and the Department for Environment, Food and Rural Affairs (Defra), grant number: SE2621 and Biotechnology and Biological Sciences Research Council (BBSRC) through projects BBS/E/I/00007030, BBS/E/I/00007033, BBS/E/I/00007036 and BBS/E/I/00007037. The funders had no role in study design, data collection and analysis, decision to publish, or preparation of the manuscript.

